# Study of cell to cell transmission of SARS CoV 2 virus particle using gene network from microarray data

**DOI:** 10.1101/2020.05.26.116780

**Authors:** Anamika Basu, Anasua Sarkar, Ujjwal Maulik

## Abstract

Microarray data from SARS CoV2 patients helps to construct a gene network relating to this disease. Analysis of these genes, present in network, highlights their biological functions and related cellular pathways. With the assistance of these information, a drug(s) can be identified to treat COVID-19. A detailed analysis of the human host response to SARS CoV 2 with expression profiling by high throughput sequencing has executed with primary human lung epithelium cell lines. Clustered genes annotation and gene network construction are performed with the help of String database. Among four cluster of genes from microarray data, cluster 1 is identified as basal cells with p= 1.37e^−05^ with 44 genes. Functional enrichment analysis of genes present in gene network has completed using String database, ToppFun online tool and NetworkAnalyst tool. For SARS CoV2 virus particles, keratin proteins, which are part of cytoskeleton structure of host cell, play a major role in cell to cell virus particle transmission. Among three types of cell- cell communication, only anchoring junction between basal cell membrane and basal lamina, is involved in this virus transmission. In this junction point, hemidesmosome structure play a vital role in virus spread from one cell to basal lamina in respiratory tract. In this protein complex structure, keratin, integrin and laminin proteins of host cell is used to promote the spread of virus infection into extracellular matrix. So, small molecular blockers of different anchoring junction proteins i.e. ITGA3, ITGA2 can provide efficient protection against this deadly virus disease. Understanding the human host response against this virus is very important to develop novel therapeutics for the treatment of SARS-CoV 2.

## I. Introduction

With the help of microarray data from a sample in a single experiment become a powerful tool for analyzing genes, responsible for a particular disease. The result of that specific condition with microarray experiment ultimately summarized in a list of genes of interest. These genes are expressed differentially in normal and diseased conditions [1]. Analysis of these genes of interest in terms of their biological functions and cellular pathways e.g. metabolic, signaling etc., helps us to identify drugs to treat this specific disease. Gene network regarding any disease means a graph with genes as vertices and edges represent relationship between genes with the help of their biological functions.

Understanding the human host response against this virus is very important to develop novel therapeutics for the treatment of SARS-CoV 2. Similar to that of other coronaviruses, this SARS-CoV 2 also causes mild to severe respiratory tract infection with very high global mortality rate [2]. Comparing differentially expressed genes in different cell lines such as primary human lung epithelium (NHBE), transformed lung alveolar (A549) cells in lung adenocarcinoma, lung biopsy cells from both normal and COVID-19 patients [3], a gene network can be constructed.

Normal human bronchial epithelial (NHBE) cells is a complex structure, composed of ciliated, non-ciliated, goblet cells and basal cells. Bronchial epithelium can be considered as a barrier against various allergens, bacteria, viruses and pollutants in between airspace and internal system of lungs and causes inflammatory response as a part of innate immunity. Immunomodulatory role of NHBE cells can be correlated with their expression of cytokines, growth factors and adhesion molecules. During this inflammatory response, leucocyte recruitment from blood increases and within bronchi by altering cell apoptosis, survival of inflammatory cells is also increased [4]. Other than cytokines, activated epithelial cells also secret leukotrienes, prostaglandins and extracellular matrix components. Thus, epithelial cells also provide an adaptive immune response against infections (viruses, bacteria and fungi), allergens and air pollutants. This is the preliminary mechanism for antigen neutralization and elimination.

All viruses employ specific mechanisms for their attachment with host cells. During virus-host interactions different classes of biomolecules are responsible for invasion and replication of viruses inside the host cells [5]. Replication of viruses is obstructed by cillia, glycolyx and immunoglobulins. Viruses enter through the epithelium layers in respiratory system by using components of cell-cell adhesion structures [6].

## II. Methodology

### i) Protein from microarray dataset analyzed and used for construction of a protein-protein network

#### a) Processing of microarray dataset

An in-depth analysis of the host response to SARS-CoV-2 with expression profiling by high throughput sequencing has executed with different cell lines such as primary human lung epithelium (NHBE), transformed lung alveolar (A549) cells and transformed lung-derived Calu-3 cells, in normal conditions and infected with SARS-CoV-2 [3] and deposited as GSE147507_RawReadCounts_Human.tsv.gz file in Series GSE147507 in GEO (Gene Expression Omnibus) [7].

Analysis of single cell RNA sequencing data with GSE147507_RawReadCounts_Human is performed using alona [8] as follows:

1. Quality filtering Uninformative genes are deleted by requiring each gene to be expressed in 1% of cells. Doublet detection is performed in this step using the Scrublet package [9].
2. Normalization Standard normalization procedure by scaling with the total read depth per cell and then multiplying with a scaling factor is used.
3. Highly variable genes (HVG) detection Highly variable genes (HVG) are discovered using a Seurat-like strategy, utilizing binning of genes according to average expression [10]. Here, HVG number is 1000 for this analysis.
4. Dimensionality reduction. Principal component (PC) analysis is performed on the HVG with the method described by Baglama and Reichel (2005) [11] with the default setting is to identify 40 PCs. 2d embedding method is completed on PCs with tDistributed Stochastic Neighbor Embedding (t-SNE) [12] with perplexity is set to 30.
5. Clustering The principal components are searched for k nearest neighbors using the BallTree algorithm. A shared nearest neighbor graph, with weights as the number of shared neighbors, is created. Cell clusters are documented from the graph with the Leiden [13] algorithm. For Leiden algorithm, cluster resolution is set to 0.6 (lesser value means larger clusters and vice versa).
6. Cell type annotation. Annotation of cell types is performed at the cluster level [14]. Gene expression in clusters is characterized by taking the median across all cells. The procedure evaluates gene expression activity of a set of marker genes and ultimately ranks the resulting cell types. Significance is calculated by figuring a one-sided Fisher’s exact test for each cell type and adjusting p-values with the Benjamini-Hochberg procedure. Thus, when the adjusted p-value is higher than 0.1, the cell type is marked with an “Unknown” annotation.
7. Differential gene expression identification This step includes all-versus-all cluster assessments; i.e., every gene is compared between every pair of clusters. Here, linear models followed by t-tests, similar to the limma R package [15] is used to generate the initial set of comparisons.

#### b) Gene network with String with clustered genes

Clustered genes are annotated and gene network is constructed by using String database [16].

### ii) Gene enrichment analysis with gene cluster present in protein-protein network

ToppFun [17], an online tool is used for functional enrichment analysis based on transcriptome, ontology, phenotype, proteome, and pharmacome annotations.

Detection of functional enrichment of forty-four gene list based on Transcriptome, Proteome, Regulome (TFBS and miRNA), Ontologies (GO, Pathway), Phenotype (human disease and mouse phenotype), Pharmacome (Drug-Gene associations), literature co-citation, and other features is performed.

## III. Result

### i) Protein from microarray dataset analyzed and used for construction of a protein-protein network

#### a) Processing of microarray dataset

With GSE147507_RawReadCounts_Human.tsv.gz file by using alona [8], a web server, is used to process and analyze the raw data with MADs value is considered as 3. MADs represents the number of median absolute deviations below the median of two quality metrics for which a cell is removed. UMAP (Uniform Manifold Approximation and Projection) algorithm to be used for 2d projection of the data.

Four cluster of proteins are formed and among them, three clusters are found with at least ten cells. 2d overview using reduction with UMAP for meta data as clusters, shows four cluster of proteins. Among them cell type annotation result shows that cluster 1 is identified as basal cells with p= 1.37e^−05^ with 44 genes (shown in Figure 1).

**Figure 1.**
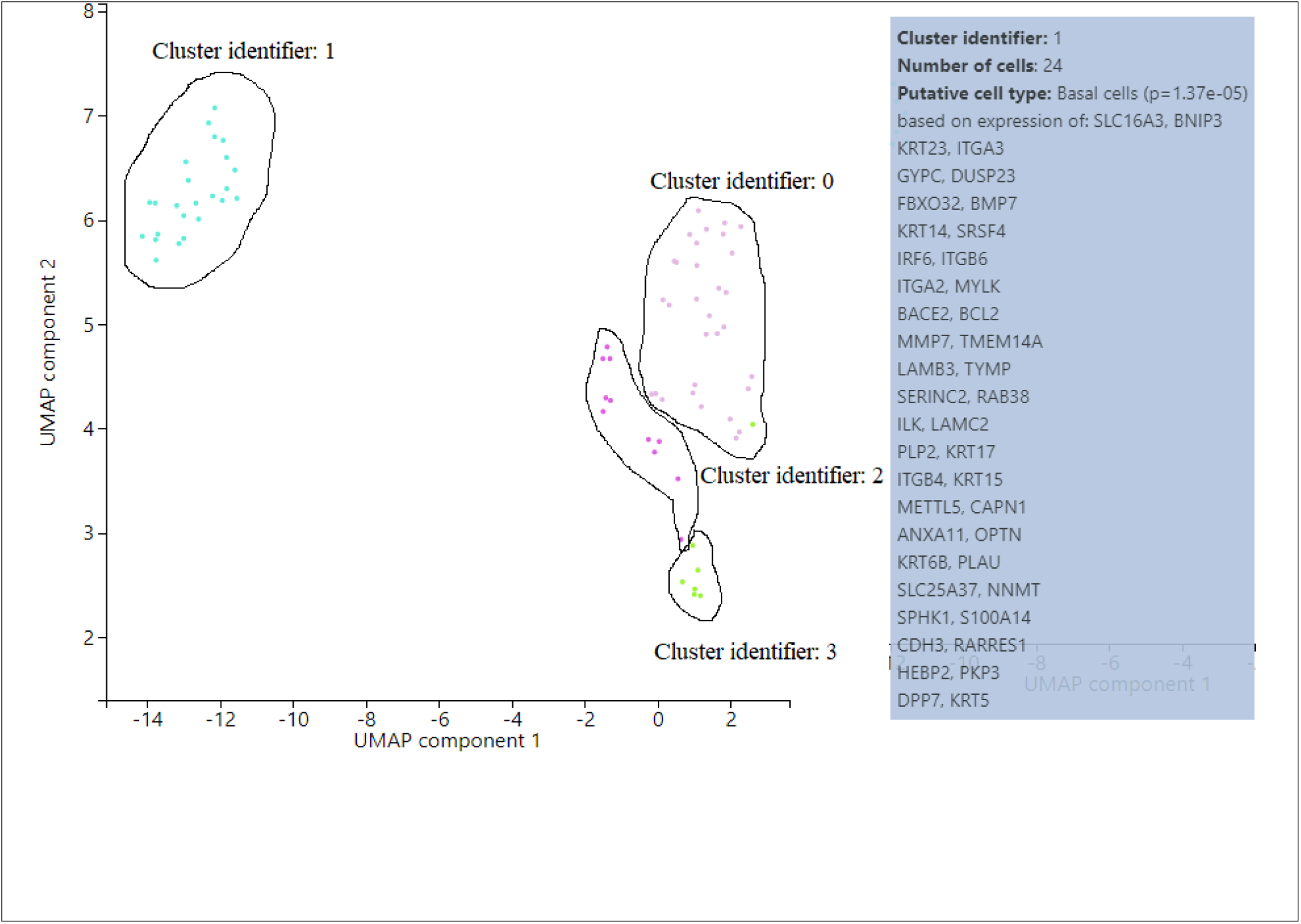
2D projection of microarray data showing four gene clusters

Twenty top expressed genes from cluster 1 for putative basal cells, are shown in Figure 2(a). Several genes e.g. KRT6A, KRT5, KRT14, KRT19 belong to keratin gene family. Box plots for two F-box proteins e.g. FBXO3 and FBXO32 are shown in Figure 2(b) and 2(c) respectively.

**Figure 2.**
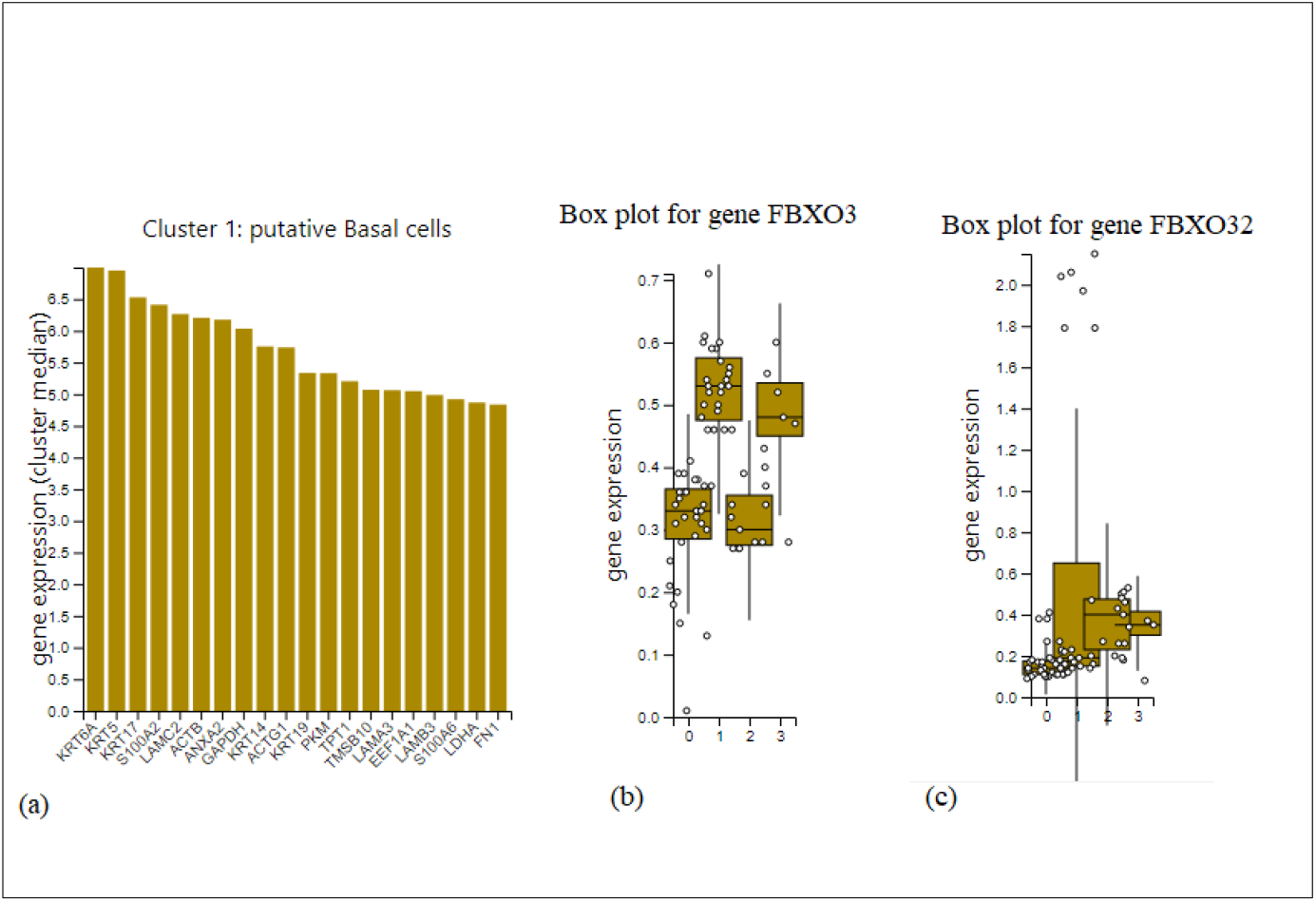
Top expressed genes in cluster 1 and box plots for two genes

#### b) Gene network with String with clustered genes

Forty-four genes in cluster 1 are annotated (Table 1) using String database [16] and used for gene network construction. Detailed annotation is presented in Supplementary table1.

**Table 1.**
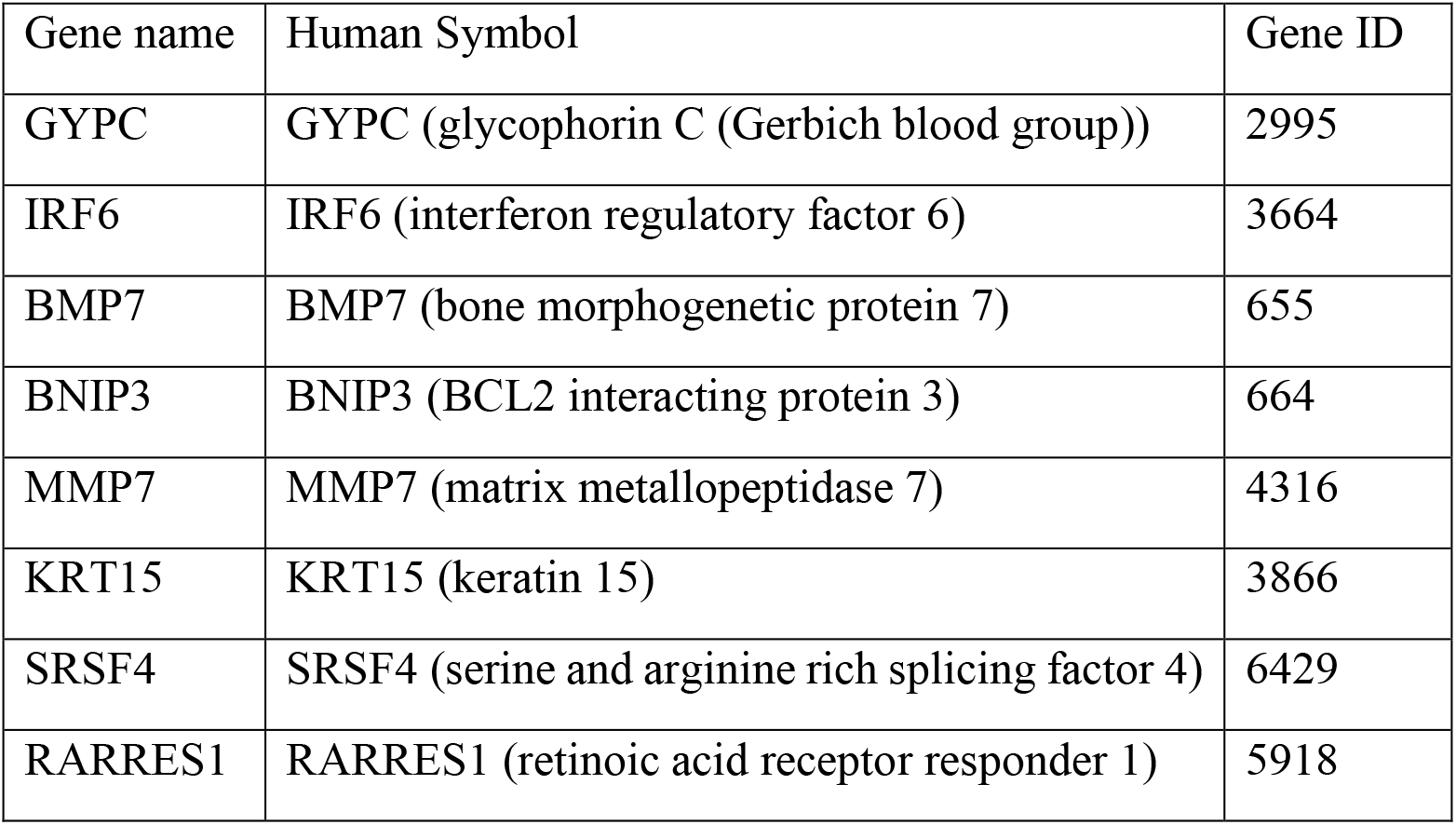
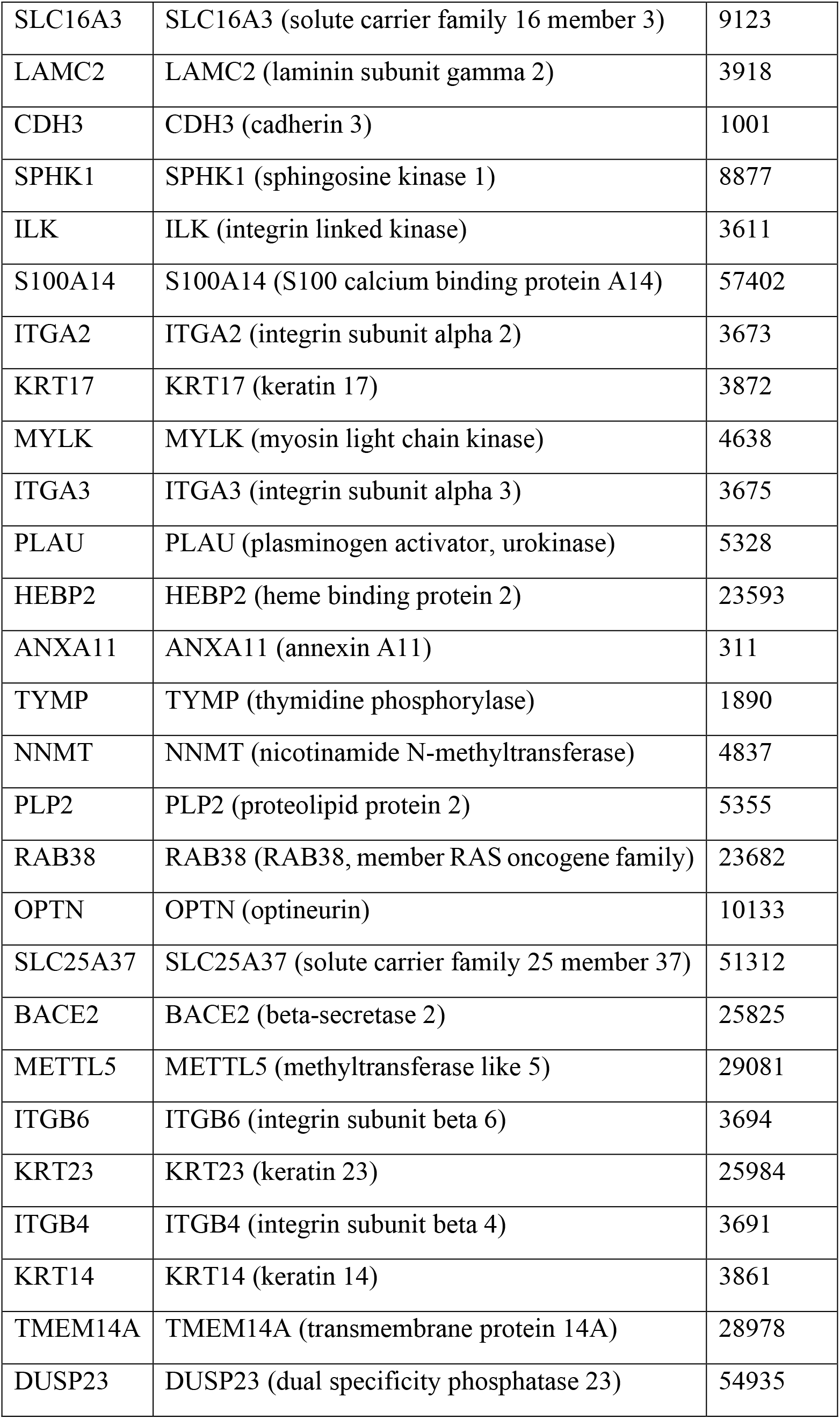
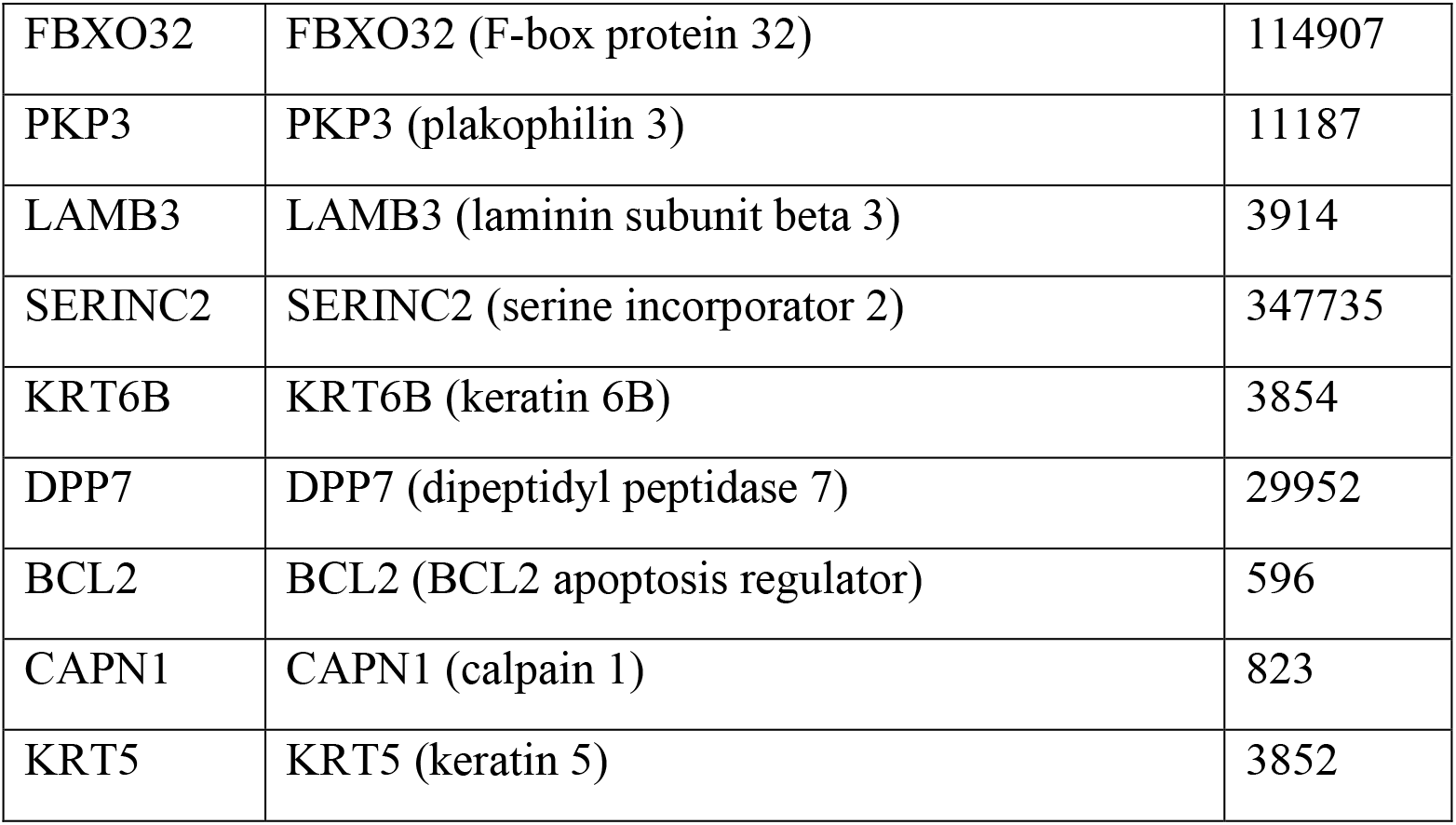
Genes present in cluster 1

Gene network construction using String database [16] with forty-four genes, obtained from cluster 1, for species *Homo sapiens*.

In this network (Figure 3) number of nodes are 44, number of edges are 59 with average node degree is 2.68. Here, avg. local clustering coefficient is 0.372 with expected number of edges is 8. PPI enrichment p-value for this network is< 1.0e-16.

**Figure 3.**
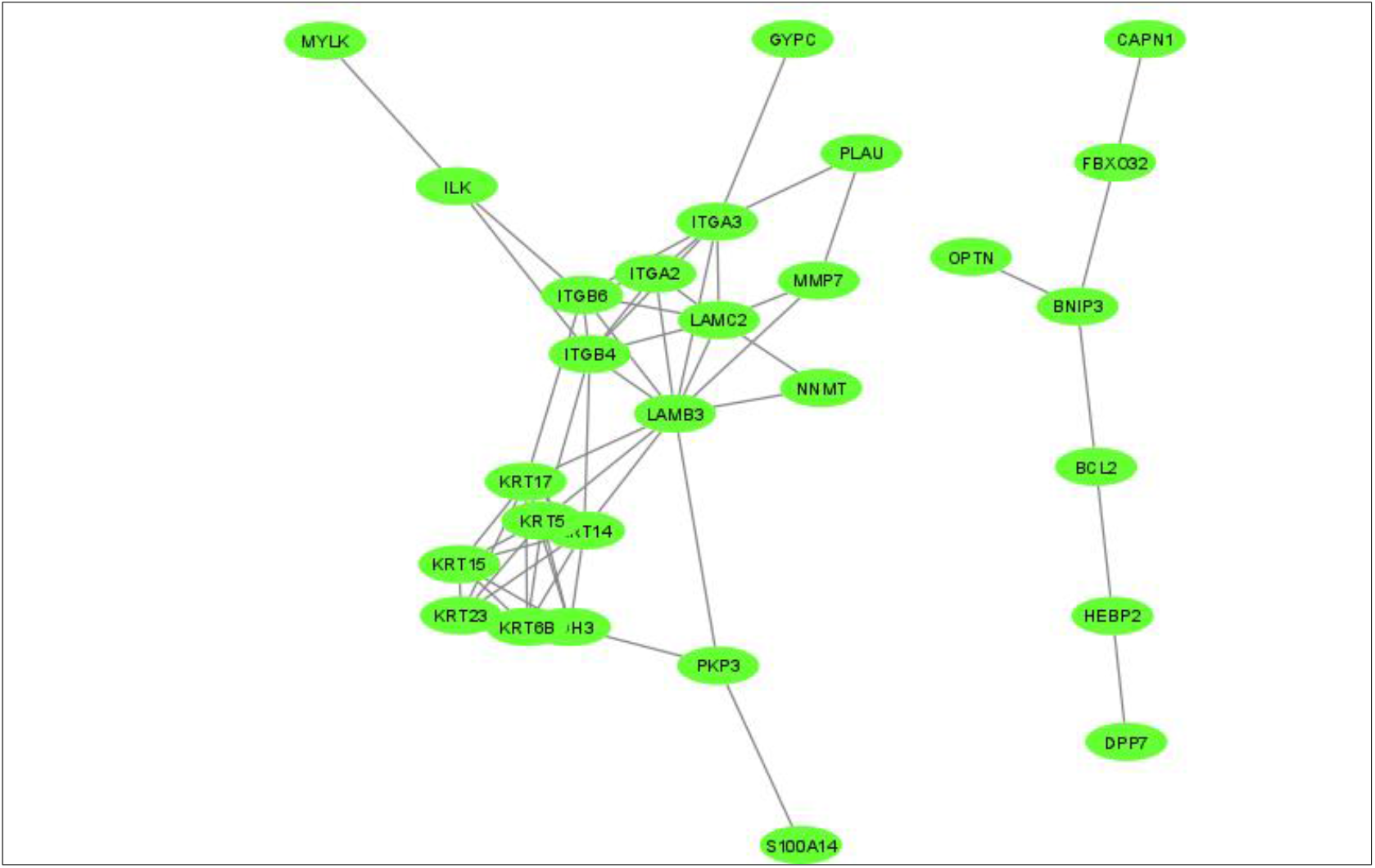
Gene network from microarray data

#### c) Functional enrichment of protein present in gene network using String database

Functional enrichment in gene network according to String database [16] considering the biological process show that 11 genes are related with cell adhesion process, 14 genes are related to skin development and 13 genes are involved in epidermis development process. 12 genes are related with epithelium development. Interestingly, 10 genes are involved in keratinocyte differentiation biological process.

Three molecular functions such as cell adhesion molecule binding (6 genes), structural constituent of cytoskeleton (5 genes e.g. KRT14, KRT17, KRT5, KRT15, KRT6B) and integrin binding (4 genes such as ILK, ITGB6, ITGA2 and ITGA3) are important in function enrichment analysis. 6 genes are present in both anchoring junction and intermediate filament whereas in integrin complex 4 genes are present. ILK, ITGB6, ITGA2, GYPC, PKP3, CDH3 are present in anchoring junction. Genes with molecular functions, known as cell adhesion molecule binding, are GYPC, ITGA3, ITGA2, ITGB6, ILK, PKP3. Among forty-four genes, present in network, only 7 genes i.e. NNMT, ITGA3, ITGA2, CDH3, BCL2, BNIP3 and FBXO32, show response to drug. According to KEGG pathway analysis, 9 genes that are MYLK, IKL, LAMC2, ITGB6, ITGB4, ITGA2, LAMB3, ITGA3 and BCL2 are present in hsa04510 pathway, which is known as focal adhesion pathway. Similarly, 7 genes namely, LAMC2, ITGB6, ITGA2, ITGB4, ITGA3, LAMB3, BCL2, are participated in PI3K-Akt signaling pathway (KEGG pathway ID hsa 04151).

Considering reactome pathway (Reactome ID HAS-446107), 5 genes i.e. LAMC2, ITGB4, LAMB3, KRT14 and KRT5, are present in type I hemidesmosome assembly. Hemidesmosomes (HDs) are specialized multiprotein junctional complexes, present as cell junction organization in cell- cell communication system. HDs connect the keratin cytoskeleton of epithelial cells to the extracellular matrix to maintain the tissue structure and integrity [18]. These complexes mediate adhesion of epithelial cells to the underlying basement membrane in different types of epithelia [19].

Hemidesmosomes are one type of anchoring junctions, present in mammalian cell junctions. Anchoring junctions form a mechanical connection between cells. Different cell junctions are involved in virus-host interactions in different viral diseases. For example, human papilloma virus induces some changes in the organizational structures of anchoring junctions. Proteins (LAMC2, LAMB3, ITBB6, ITGA2, ITGA3, ITGB4) present in anchoring junctions of epithelial cells, are related with KEGG pathway of human papillomavirus infection (KEGG pathway ID hsa 05165). Rotavirus and hepatitis C virus (HCV) spread through tight junctions, which is another type of cell junction of human body. Similarly, human immune deficiency virus (HIV) penetrates through the epithelial barrier by damaging gap junctions [20].

The precise entry point of virus in cellular entry can be predicted as target for possible therapeutics against viral infection. Blockers of theses anchoring junction proteins i.e. ITGA3, ITGA2 can provide protection against SARS-CoV 2 infection. Six genes i.e. LAMC2, ITGA3, ITGB4, LAMB3, KRT14 and KRT5, in this network, are related with epidermolysis bullosa (UniProt Keyword Id KW-0263). Epidermolysis bullosa is a group of inherited connective tissue diseases, characterized by multiple blisters on skin and respiratory and gastrointestinal tract mucosa. But till now no evidence of COVID-19 infection with epidermolysis bullosa patient has been reported.

Six intermediate filament proteins i.e. KRT14, KRT17, KRT5, KRT15, KRT6B, KRT23 are proteins which are primodial components of the cytoskeleton and the nuclear envelop, with filamentous structures [21]. For many viral infections intermediate filaments play novel role during infection. For example, parovirus minute virus of mice (MVM) intermediate filament has been altered. Vimentin, which is an example of intermediate filament, participates in cell adhesion, related with several viral diseases such as SARS-CoV, dengue, African swine fever etc [22]. Different pathogens e.g. herpes simplex virus, human papilloma virus type 16, adenovirus, rotavirus, rhinovirus, can cause keratin intermediate filament network disruption. In HPV infection, keratins are hyperphosphorylated and ubiquitinylated causing keratin network collapse, resulting increased release of viral particles for further infection. Thicker keratin filament and disruption in keratin network occur in herpes simplex virus type 2 (HSV-2) infection. Keratin network disruption is also related with cell lysis of host cell and release of mature virus particles during human rhinovirus infection [23].

#### d) Gene enrichment analysis with gene cluster present in protein-protein network using ToppFun tool

Probability density function is the p value method for gene list enrichment analysis with ToppFun [17], online tool. ToppFun analysis result shows that among forty-four genes, 10 proteins from 10 genes (KRT23, KRT5, KRT6B, SERINC2, KRT14, KRT15, KRT17, SPHK1, PKP3 and CAPN1) have transferase activity for transferring acyl groups (GO:0016746). Except SERINC2 and SPHK1, other eight proteins are related with glutamine gamma-glutamyl transferase activity (GO:0003810). 9 genes are present as cellular component of anchoring junction. These genes are PLAU, ITGA2, ILK, ITGA3, CDH, ITGB4, ITGB6, PKP3 and CAPN1. Genes related with different pathways are summarized in Table 2.

**Table 2.** Gene enrichment analysis according to their molecular functions

|  | ID | Name | pValue | Bonferroni | Genes from Input | Genes in Annotation |
| --- | --- | --- | --- | --- | --- | --- |
| 1 | GO:0003810 | protein-glutamine gamma-glutamyltransferase activity | 3.685E-10 | 9.691E-8 | 8 | 125 |
| 2 | GO:0016755 | transferase activity, transferring amino-acyl groups | 6.431E-10 | 1.691E-7 | 8 | 134 |
| 3 | GO:0016746 | transferase activity, transferring acyl groups | 2.146E-8 | 5.644E-6 | 10 | 408 |
| 4 | GO:0005200 | structural constituent of cytoskeleton | 6.035E-6 | 1.587E-3 | 5 | 114 |
| 5 | GO:0005178 | integrin binding | 2.009E-5 | 5.284E-3 | 5 | 146 |
| 6 | GO:0050839 | cell adhesion molecule binding | 1.726E-4 | 4.540E-2 | 7 | 525 |
| 7 | GO:0005198 | structural molecule activity | 3.381E-4 | 8.893E-2 | 8 | 779 |

#### e) Gene annotation by NetworkAnalyst

By using NetworkAnalyst tool [24], a network is drawn (Figure 4) for gene enrichment analysis on the basis of KEGG pathway. Focal adhesion pathway has the highest p value (7.957E^−9^). The genes which are present in focal adhesion pathway are LAMB3, LAMC2, BCL2, ITGA2, ILK, ITGA3, MYLK, ITGB4 and ITGB6. According to Wilkipathways at the cell-extracellular matrix contact points, a specialized structure, known as focal adhesion is formed. Here actin filament bundles are attached with integrin family protein. Similarly, proteins from genes LAMB3, LAMC2, BCL2, ITGA2, ITGA3, ITGB4 and ITGB6, are present in PI3K-Akt signaling pathway (Table 3). During viral infection, this pathway is an important mechanism through which virus influences various cell functions of the host. For example, apoptosis is slowed down and virus replication process is prolonged using this pathway [25]. Thus, virus particles are hijacking the PI3K-Akt signaling pathway of the host cell.

**Figure 4.**
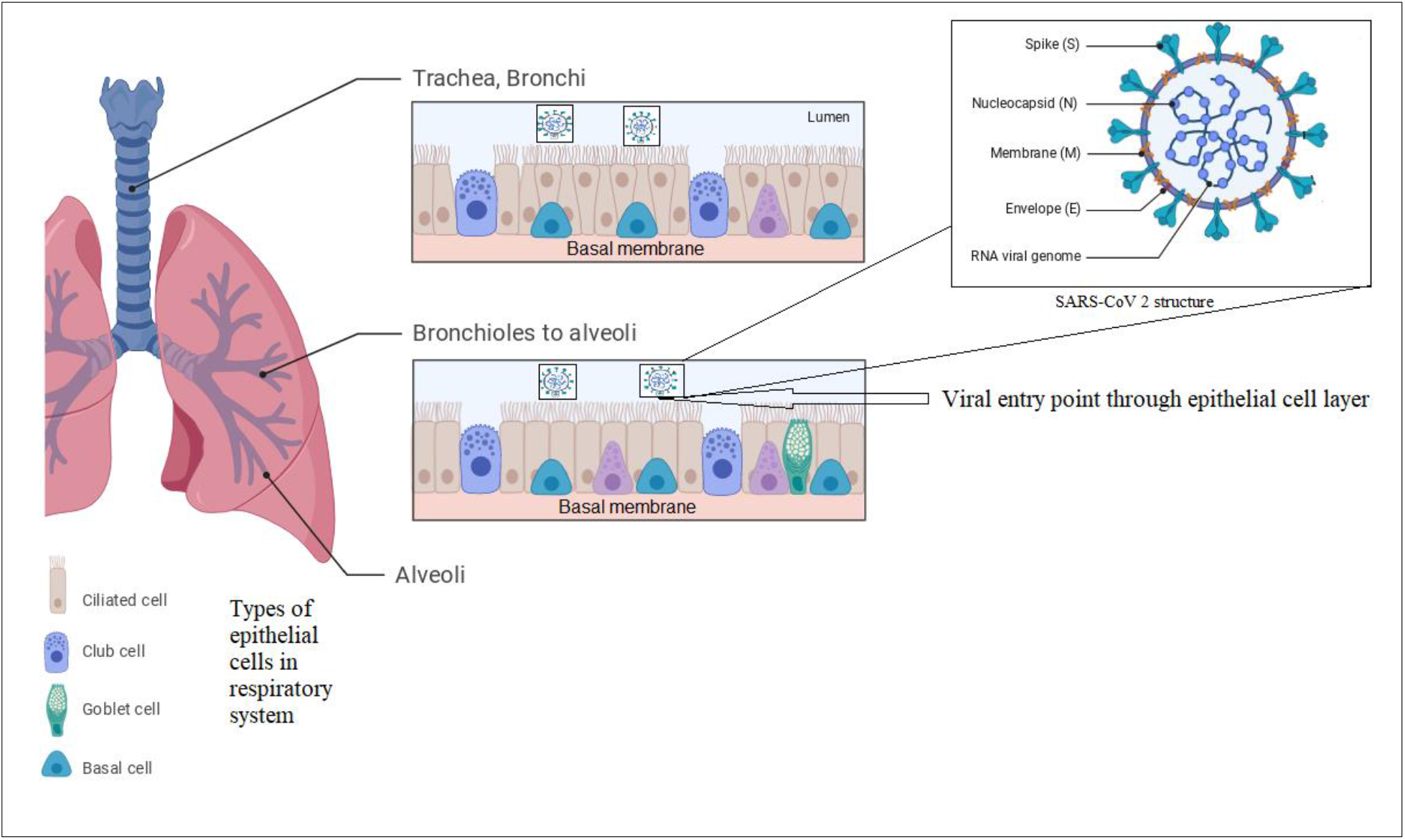
Virus entry point through respiratory system

**Table 3.** Biological pathways related with cluster 1 genes

| Name of Pathway | Total genes present in the pathway | Expected frequency | No. of Hits | PValue | FDR |
| --- | --- | --- | --- | --- | --- |
| Focal adhesion | 199 | 0.72 | 9 | 1.86e-08 | 5.92e-06 |
| ECM-receptor interaction | 82 | 0.297 | 6 | 3.68e-07 | 5.86e-05 |
| Small cell lung cancer | 93 | 0.337 | 5 | 1.78e-05 | 0.00189 |
| Estrogen signaling pathway | 138 | 0.499 | 5 | 0.000119 | 0.00758 |
| Arrhythmogenic right ventricular cardiomyopathy (ARVC) | 72 | 0.261 | 4 | 0.000119 | 0.00758 |
| PI3K-Akt signaling pathway | 354 | 1.28 | 7 | 0.000203 | 0.0103 |
| Hypertrophic cardiomyopathy (HCM) | 85 | 0.308 | 4 | 0.000227 | 0.0103 |
| Dilated cardiomyopathy | 91 | 0.329 | 4 | 0.000295 | 0.0117 |
| Regulation of actin cytoskeleton | 214 | 0.774 | 5 | 0.000903 | 0.0319 |
| Pathways in cancer | 530 | 1.92 | 5 | 0.0392 | 0.999 |

**Figure 5.**
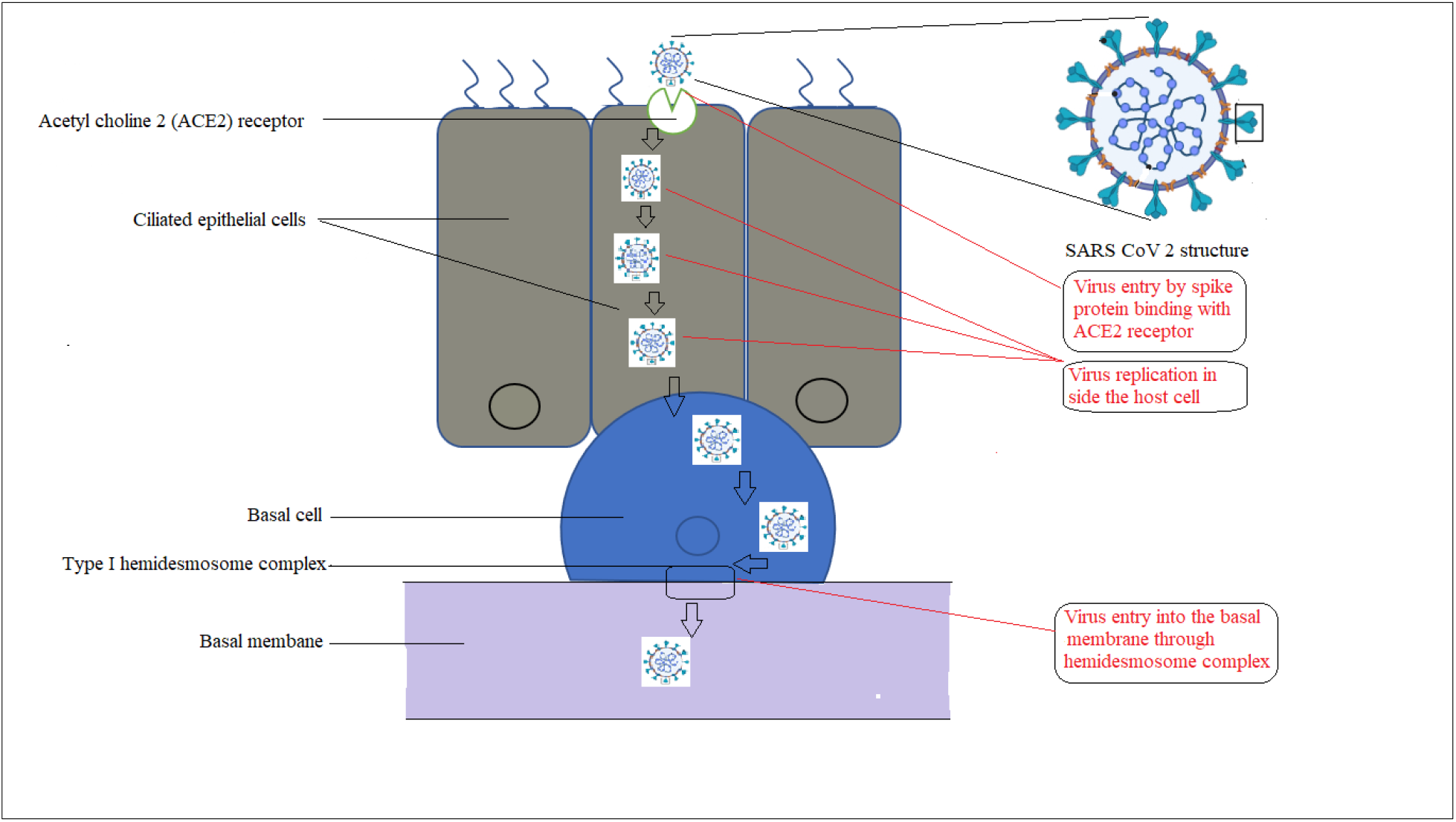
Diagramatic representation of virus entry and cell to cell transmission

**Figure 6.**
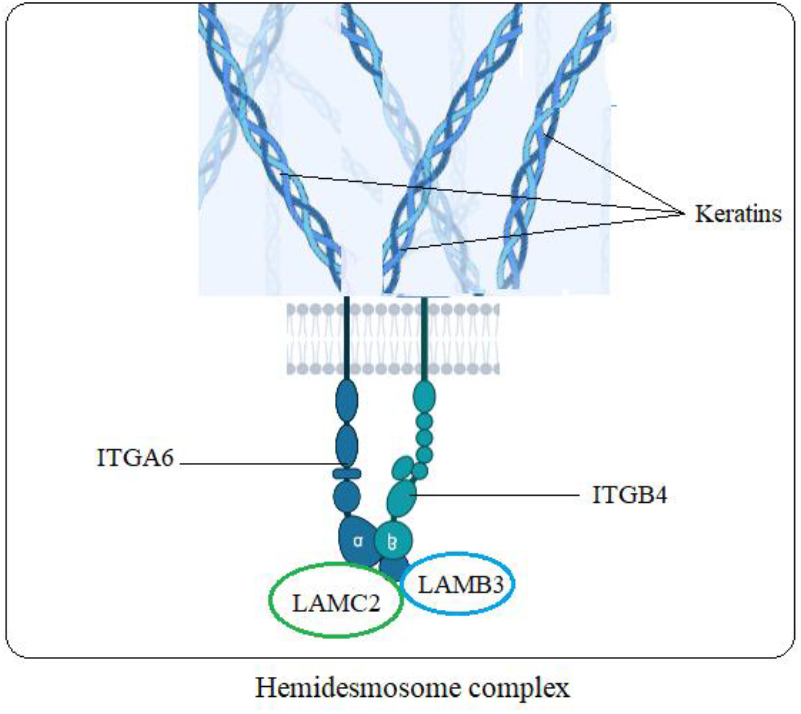
Structure of hemidesmosome complex

**Figure 7.**
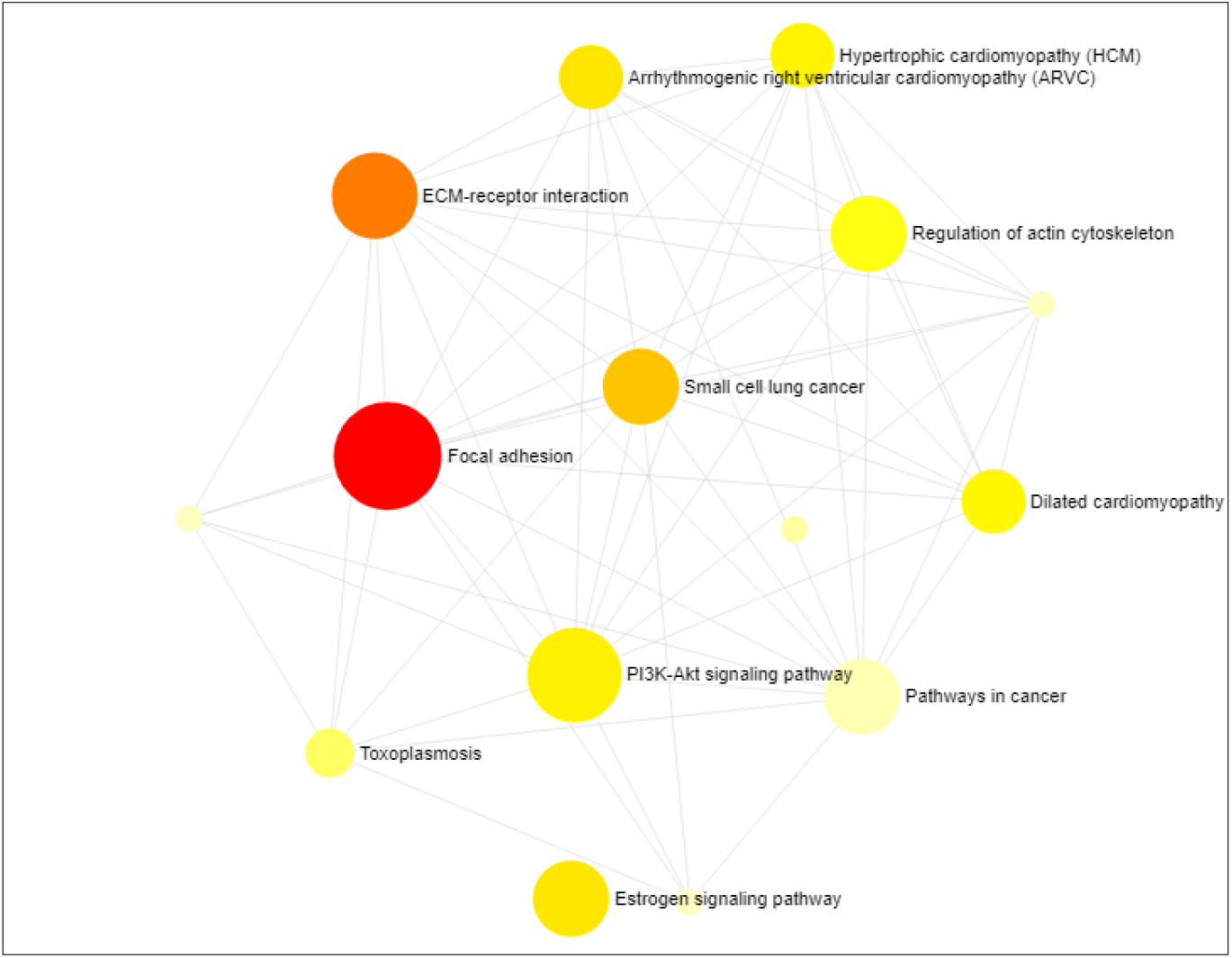
Enrichment analysis of cluster 1 genes

In earlier studies Mizutani et al, 2004, have proved that SARS-CoV coronavirus infection, PI3K/Akt pathway is activated in virus-infected cells [26]. Similarly, for Middle East Respiratory syndrome coronavirus (MERS-CoV), kinase inhibitors targeting PI3K/Akt signaling pathway, are considered as potential antiviral agents in MERS-CoV infection [27].

## IV. Discussion

In respiratory tract infection, epithelial cell layer can act as a first line of defense in innate immune response. Among different types of cells, ciliated cells which are attached with basal membrane through basal cells, can be considered as the entry point of SARS CoV2 viral infection. This virus particles enter into the human host cells through the binding to the cell surface receptor protein angiotensin converting enzyme 2 (ACE2) [28]. Surface binding spike protein of virus mediates its entry by binding with host membrane receptor protein. After entry inside the host cell in human, virus particles move via endosome formation and finally fuse with lysosomal membranes. Then, envelop protein of virus particles disrupt and their genetic material, RNA released into the host cell. By using replication machinery of host cell, more virus particles are generated.

Virus particles can spread through the host body by two distinct methods. First method is diffusion through extracellular matrix and the second one is by using direct cell to cell contact [29]. Viruses which are transmitted across tight cell to cell contacts, are reached to neighborhood cells and extracellular matrix via host cell surface receptors. Thus, their binding efficiency and entry inside the host cell are increased manifolds. For SARS CoV2 virus particles, keratin proteins, which are part of cytoskeleton structure of host cell, play a major role in cell to cell transmission. Basal cell of epithelial cell layer is attached with basal membrane through cell to cell junction points. Among three junction points in cell to cell contact in human body, in SARS CoV2 infection, only anchoring junction between basal cell membrane and basal lamina, is involved in virus transmission. In this junction point, hemidesmosome structure play a vital role in virus spread from one cell to basal lamina in respiratory tract. Integrin proteins which binds to laminin proteins, are present in extracellular matrix, form hemidesmosome structure. Thus, focal adhesion pathway is affected in this cell to cell virus transmission. Although HCV and HIV viruses, transmit through tight junctions and gap junctions but SARS CoV2 virus transmits by using anchoring junction points. Here, keratin, integrin and laminin proteins of host cell are used to promote the spread of virus infection into extracellular matrix. The precise protein receptor i.e. ACE2 can be used as target for possible therapeutics in SARS CoV2 treatment and at the time small molecular blockers of different anchoring junction proteins i.e. ITGA3, ITGA2 can provide efficient protection against this deadly virus disease.

