## Supplementary table 1 for "Study of cell to cell transmission of SARS CoV 2 virus particle using gene network from microarray data"

| Sl no. | Gene name | Gene annotation |
| --- | --- | --- |
| 1 | GYPC | Glycophorin-C; This protein is a minor sialoglycoprotein in human erythrocyte membranes. The blood group Gerbich antigens and receptors for Plasmodium falciparum merozoites are most likely located within the extracellular domain. Glycophorin-C plays an important role in regulating the stability of red cells |
| 2 | IRF6 | Interferon regulatory factor 6; Probable DNA-binding transcriptional activator. Key determinant of the keratinocyte proliferation-differentiation switch involved in appropriate epidermal development (By similarity). Plays a role in regulating mammary epithelial cell proliferation (By similarity). May regulate WDR65 transcription (By similarity); Belongs to the IRF family |
| 3 | BMP7 | Bone morphogenetic protein 7; Induces cartilage and bone formation. May be the osteoinductive factor responsible for the phenomenon of epithelial osteogenesis. Plays a role in calcium regulation and bone homeostasis; Promotes brown adipocyte differentiation by P38 MAPK pathway-mediated activation of target genes, including members of the SOX family of transcription factors |
| 4 | BNIP3 | BCL2/adenovirus E1B 19 kDa protein-interacting protein 3; Apoptosis-inducing protein that can overcome BCL2 suppression. May play a role in repartitioning calcium between the two major intracellular calcium stores in association with BCL2. Involved in mitochondrial quality control via its interaction with SPATA18/MIEAP: in response to mitochondrial damage, participates in mitochondrial protein catabolic process (also named MALM) leading to the degradation of damaged proteins inside mitochondria. The physical interaction of SPATA18/MIEAP, BNIP3 and BNIP3L/NIX at the mitochondrial outer membrane regulates the opening of a pore in the mitochondrial double membrane in order to mediate the translocation of lysosomal proteins from the cytoplasm to the mitochondrial matrix |
| 5 | MMP7 | Matrilysin; Degrades casein, gelatins of types I, III, IV, and V, and fibronectin. Activates procollagenase; M10 matrix metallopeptidases |
| 6 | KRT15 | Keratin, type I cytoskeletal 15; Keratins, type I |
| 7 | SRSF4 | Serine/arginine-rich splicing factor 4; Plays a role in alternative splice site selection during pre-mRNA splicing. Represses the splicing of MAPT/Tau exon 10; RNA binding motif containing |
| 8 | RARRES1 | Retinoic acid receptor responder protein 1; Inhibitor of the cytoplasmic carboxypeptidase AGBL2, may regulate the alpha-tubulin tyrosination cycle |
| 9 | SLC16A3 | Monocarboxylate transporter 4; Proton-linked monocarboxylate transporter. Catalyzes the rapid transport across the plasma membrane of many monocarboxylates such as lactate, pyruvate, branched-chain oxo acids derived from leucine, valine and isoleucine, and the ketone bodies acetoacetate, beta-hydroxybutyrate and acetate (By similarity); Belongs to the major facilitator superfamily |
| 10 | LAMC2 | Laminin subunit gamma-2; Binding to cells via a high affinity receptor, laminin is thought to mediate the attachment, migration and organization of cells into tissues during embryonic development by interacting with other extracellular matrix components. Ladsin exerts cell- scattering activity toward a wide variety of cells, including epithelial, endothelial, and fibroblastic cells |
| 11 | CDH3 | Cadherin-3; Cadherins are calcium-dependent cell adhesion proteins. They preferentially interact with themselves in a homophilic manner in connecting cells; cadherins may thus contribute to the sorting of heterogeneous cell types |
| 12 | SPHK1 | Sphingosine kinase 1; Catalyzes the phosphorylation of sphingosine to form sphingosine 1-phosphate (SPP), a lipid mediator with both intra- and extracellular functions. Also acts on D-erythro- sphingosine and to a lesser extent sphinganine, but not other lipids, such as D,L-threo-dihydrosphingosine, N,N- dimethylsphingosine, diacylglycerol, ceramide, or phosphatidylinositol |
| 13 | ILK | Integrin-linked protein kinase; Receptor-proximal protein kinase regulating integrin- mediated signal transduction. May act as a mediator of inside-out integrin signaling. Focal adhesion protein part of the complex ILK-PINCH. This complex is considered to be one of the convergence points of integrin- and growth factor-signaling pathway. Could be implicated in mediating cell architecture, adhesion to integrin substrates and anchorage- dependent growth in epithelial cells. Phosphorylates beta-1 and beta-3 integrin subunit on serine and threonine residues, but also AKT1 and GSK3B |
| 14 | S100A14 | Protein S100-A14; Modulates P53/TP53 protein levels, and thereby plays a role in the regulation of cell survival and apoptosis. Depending on the context, it can promote cell proliferation or apoptosis. Plays a role in the regulation of cell migration by modulating the levels of MMP2, a matrix protease that is under transcriptional control of P53/TP53. |
| 15 | ITGA2 | Integrin alpha-2; Integrin alpha-2/beta-1 is a receptor for laminin, collagen, collagen C-propeptides, fibronectin and E-cadherin. It recognizes the proline-hydroxylated sequence G-F-P-G-E-R in collagen. It is responsible for adhesion of platelets and other cells to collagens, modulation of collagen and collagenase gene expression, force generation and organization of newly synthesized extracellular matrix |
| 16 | KRT17 | Keratin, type I cytoskeletal 17; Type I keratin involved in the formation and maintenance of various skin appendages, specifically in determining shape and orientation of hair (By similarity). Required for the correct growth of hair follicles, in particular for the persistence of the anagen (growth) state (By similarity). Modulates the function of TNF-alpha in the specific context of hair cycling. Regulates protein synthesis and epithelial cell growth through binding to the adapter protein SFN and by stimulating Akt/mTOR pathway (By similarity). Involved in tissue repair. May be a marker of basal cell differentiation in complex epithelia and therefore indicative of a certain type of epithelial stem cells |
| 17 | MYLK | Myosin light chain kinase, smooth muscle; Calcium/calmodulin-dependent myosin light chain kinase implicated in smooth muscle contraction via phosphorylation of myosin light chains (MLC). Also regulates actin-myosin interaction through a non-kinase activity. Phosphorylates PTK2B/PYK2 and myosin light-chains. Involved in the inflammatory response (e.g. apoptosis, vascular permeability, leukocyte diapedesis), cell motility and morphology, airway hyperreactivity and other activities relevant to asthma. Required for tonic airway smooth muscle contraction that is necessary for physiological and asthmatic airway resistance |
| 18 | ITGA3 | Integrin alpha-3; Integrin alpha-3/beta-1 is a receptor for fibronectin, laminin, collagen, epiligrin, thrombospondin and CSPG4. Integrin alpha-3/beta-1 provides a docking site for FAP (seprase) at invadopodia plasma membranes in a collagen-dependent manner and hence may participate in the adhesion, formation of invadopodia and matrix degradation processes, promoting cell invasion. |
| 19 | PLAU | Urokinase-type plasminogen activator; Specifically cleaves the zymogen plasminogen to form the active enzyme plasmin |
| 20 | HEBP2 | Heme-binding protein 2; Can promote mitochondrial permeability transition and facilitate necrotic cell death under different types of stress conditions |
| 21 | ANXA11 | Annexin A11; Binds specifically to calcyclin in a calcium-dependent manner (By similarity). Required for midbody formation and completion of the terminal phase of cytokinesis; Belongs to the annexin family |
| 22 | TYMP | Thymidine phosphorylase; May have a role in maintaining the integrity of the blood vessels. Has growth promoting activity on endothelial cells, angiogenic activity in vivo and chemotactic activity on endothelial cells |
| 23 | NNMT | Nicotinamide N-methyltransferase; Catalyzes the N-methylation of nicotinamide and other pyridines to form pyridinium ions. This activity is important for biotransformation of many drugs and xenobiotic compounds; |
| 24 | PLP2 | Proteolipid protein 2; May play a role in cell differentiation in the intestinal epithelium |
| 25 | RAB38 | Ras-related protein Rab-38; May be involved in melanosomal transport and docking. Involved in the proper sorting of TYRP1. Involved in peripheral melanosomal distribution of TYRP1 in melanocytes; the function, which probably is implicating vesicle-trafficking, includes cooperation with ANKRD27 and VAMP7 (By similarity). Plays a role in the maturation of phagosomes that engulf pathogens, such as S.aureus and M.tuberculosis. |
| 26 | OPTN | Optineurin; Plays an important role in the maintenance of the Golgi complex, in membrane trafficking, in exocytosis, through its interaction with myosin VI and Rab8. Links myosin VI to the Golgi complex and plays an important role in Golgi ribbon formation. Plays a role in the activation of innate immune response during viral infection. Mechanistically, recruits TBK1 at the Golgi apparatus, promoting its trans-phosphorylation after RLR or TLR3 stimulation. In turn, activated TBK1 phosphorylates its downstream partner IRF3 to produce IFN-beta. |
| 27 | SLC25A37 | Mitoferrin-1; Mitochondrial iron transporter that specifically mediates iron uptake in developing erythroid cells, thereby playing an essential role in heme biosynthesis. The iron delivered into the mitochondria, presumably as Fe(2+), is then probably delivered to ferrochelatase to catalyze Fe(2+) incorporation into protoprophyrin IX to make heme (By similarity); Solute carriers |
| 28 | BACE2 | Beta-secretase 2; Responsible for the proteolytic processing of the amyloid precursor protein (APP). Cleaves APP, between residues 690 and 691, leading to the generation and extracellular release of beta-cleaved soluble APP, and a corresponding cell-associated C- terminal fragment which is later released by gamma-secretase. It has also been shown that it can cleave APP between residues 671 and 672 |
| 29 | METTL5 | Methyltransferase-like protein 5; Probable methyltransferase |
| 30 | ITGB6 | Integrin beta-6; Integrin alpha-V/beta-6 is a receptor for fibronectin and cytotactin. It recognizes the sequence R-G-D in its ligands. Internalisation of integrin alpha-V/beta-6 via clathrin-mediated endocytosis promotes carcinoma cell invasion. ITGAV:ITGB6 acts as a receptor for fibrillin-1 (FBN1) and mediates R-G-D-dependent cell adhesion to FBN1 |
| 31 | KRT23 | Keratin, type I cytoskeletal 23; Keratins, type I |
| 32 | ITGB4 | Integrin beta-4; Integrin alpha-6/beta-4 is a receptor for laminin. Plays a critical structural role in the hemidesmosome of epithelial cells. Is required for the regulation of keratinocyte polarity and motility. ITGA6:ITGB4 binds to NRG1 (via EGF domain) and this binding is essential for NRG1-ERBB signaling. ITGA6:ITGB4 binds to IGF1 and this binding is essential for IGF1 signaling |
| 33 | KRT14 | Keratin, type I cytoskeletal 14; The nonhelical tail domain is involved in promoting KRT5-KRT14 filaments to self-organize into large bundles and enhances the mechanical properties involved in resilience of keratin intermediate filaments in vitro |
| 34 | TMEM14A | Transmembrane protein 14A; Inhibits apoptosis via negative regulation of the mitochondrial outer membrane permeabilization involved in apoptotic signaling pathway; Belongs to the TMEM14 family |
| 35 | DUSP23 | Dual specificity protein phosphatase 23; Protein phosphatase that mediates dephosphorylation of proteins phosphorylated on Tyr and Ser/Thr residues. In vitro, it can dephosphorylate p44-ERK1 (MAPK3) but not p54 SAPK-beta (MAPK10) in vitro. Able to enhance activation of JNK and p38 (MAPK14); Atypical dual specificity phosphatases |
| 36 | FBXO32 | F-box only protein 32; Substrate recognition component of a SCF (SKP1-CUL1-F- box protein) E3 ubiquitin-protein ligase complex which mediates the ubiquitination and subsequent proteasomal degradation of target proteins. Probably recognizes and binds to phosphorylated target proteins during skeletal muscle atrophy. Recognizes TERF1; F-boxes other |
| 37 | PKP3 | Plakophilin-3; May play a role in junctional plaques; Armadillo repeat containing |
| 38 | LAMB3 | Laminin subunit beta-3; Binding to cells via a high affinity receptor, laminin is thought to mediate the attachment, migration and organization of cells into tissues during embryonic development by interacting with other extracellular matrix components |
| 39 | SERINC2 | Serine incorporator 2; Belongs to the TDE1 family |
| 40 | KRT6B | Keratin, type II cytoskeletal 6B; Keratins, type II; Belongs to the intermediate filament family |
| 41 | DPP7 | Dipeptidyl peptidase 2; Plays an important role in the degradation of some oligopeptides; DASH family |
| 42 | BCL2 | Apoptosis regulator Bcl-2; Suppresses apoptosis in a variety of cell systems including factor-dependent lymphohematopoietic and neural cells. Regulates cell death by controlling the mitochondrial membrane permeability. Appears to function in a feedback loop system with caspases. Inhibits caspase activity either by preventing the release of cytochrome c from the mitochondria and/or by binding to the apoptosis-activating factor (APAF-1). May attenuate inflammation by impairing NLRP1-inflammasome activation, hence CASP1 activation and IL1B release; BCL2 family |
| 43 | CAPN1 | Calpain-1 catalytic subunit; Calcium-regulated non-lysosomal thiol-protease which catalyzes limited proteolysis of substrates involved in cytoskeletal remodeling and signal transduction; Belongs to the peptidase C2 family |
| 44 | KRT5 | Keratin, type II cytoskeletal 5; Keratins, type II |
